# *k*-Nearest Common Leaves algorithm for phylogenetic tree completion

**DOI:** 10.64898/2026.04.02.716144

**Authors:** Aleksandr Koshkarov, Nadia Tahiri

## Abstract

Phylogenetic trees represent the evolutionary histories of taxa and support tasks such as clustering and Tree of Life reconstruction. Many established comparison methods, including the Robinson-Foulds (RF) distance, assume identical taxon sets. A methodological gap remains for trees with distinct but overlapping taxa. Existing approaches either prune non-common leaves, which can discard information, or complete both trees such that they share the same taxa. Completion is more comprehensive, but current methods typically ignore branch lengths, which are essential for identifying evolutionary patterns. This paper introduces *k*-Nearest Common Leaves (*k*-NCL), an algorithm for completing rooted phylogenetic trees defined on different but overlapping taxa. The method uses branch lengths and topological characteristics and does not rely on a specific distance measure. The *k*-NCL algorithm is designed to preserve evolutionary relationships in the trees under comparison. The running time is *O*(*n*^2^), where *n* is the size of the union of the two leaf sets. Additional properties include preservation of original distances and topology, symmetry, and uniqueness of the completion. Implemented in Python, *k*-NCL is evaluated on biological datasets of amphibians, birds, mammals, and sharks. Experimental results show that RF combined with *k*-NCL improves phylogenetic tree clustering performance compared to the RF(+) tree completion approach.

**Availability and implementation:** An open-source implementation of *k*-NCL in Python and the datasets used in this study are available at https://github.com/tahiri-lab/KNCL.

## 1 Introduction

Phylogenetic trees are widely used to study the diversity of life, providing simplified representations of complex evolutionary relationships. They are utilized in fields such as comparative genomics and evolutionary biology. A common task in these areas is measuring distances between phylogenetic trees, often for purposes such as clustering trees and evaluating phylogenetic inference methods. Such distance measurements are especially informative when the trees share the same set of species. However, in many practical cases, the trees are defined on different but overlapping sets of taxa. In order to compare such trees, researchers often use methods such as pruning non-overlapping leaves or completing both trees to make them defined on the same set of taxa [1,35]. Pruning can result in the loss of valuable evolutionary information by removing unique taxa, whereas tree completion retains all taxa and allows for a more comprehensive comparison of evolutionary relationships. However, topology-only tree completion approaches may overlook important information, such as branch lengths, potentially limiting the accuracy of comparisons. This highlights the need for more advanced methods that use both topological and branch length information to provide a more accurate representation of evolutionary signals.

Practical tasks such as supertree construction [2,34,29], phylogenetic tree clustering [29,30], the assembly of the Tree of Life [11], and searches within phylogenetic databases [33,5] require the computation of distances between trees with different numbers of overlapping leaves. Approaches that can handle trees defined on different but overlapping sets of taxa include the Robinson-Foulds(-) method, which prunes both trees to a shared taxa set before computing the classical Robinson-Foulds (RF) distance [7], the generalized RF distance [17], and the vectorial tree distance [21]. The tree completion-based methods involve the addition of distinct leaves from one tree to another, thereby defining both trees on the same set of taxa, which is the union of the leaf sets of the original trees. These methods include the RF(+) approach [1,35], geodesic in the extended Billera-Holmes-Vogtmann (BHV) tree space [10,24], and several others [14,16].

The RF-based tree completion approach, detailed and expanded in various studies [7,6,15,8,1,35], uses a completion method aiming to minimize the RF distance. The RF(+) method described in [35] implements a polynomial-time dynamic programming algorithm for computing the RF(+) distance on completed trees. The RF(+) method only accounts for the topology of the trees being compared. The exact algorithm for the RF(+) problem has a complexity of *O*(*n* · *k*^2^), where *n* is the size of the union of the leaf sets of both trees under comparison, and *k* is the number of maximal subtrees unique to one input tree.

The geodesic distance in the BHV tree space [4] considers both the topology and branch lengths of the trees. Extensions of this metric to process trees with different but overlapping sets of taxa have been proposed in [10,24], introducing techniques to compute distances in an extended BHV tree space. The introduction of additional structures within the BHV tree space, such as a connection cluster, a connection space, a connection graph, and the incorporation of new leaves, which requires a transition between lower and higher dimensional orthants in the BHV space, results in an increase in computational complexity to *O*(*n*^*ℓ*+2^), where *n* is the size of the union of the leaf sets of the trees being compared and *ℓ* is the number of new leaves to be added to the tree. Although these methods allow for comprehensive distance calculations, the computational intensity required for large trees with numerous non-common leaves is a significant limitation. It should be noted that the extended BHV geodesic defines a pairwise distance via extension spaces rather than yielding a unique completed tree for each input, and therefore is not a tree completion method.

Several researchers have developed different strategies for imputing missing taxa to phylogenetic trees [18,19,22,36,37]. The approach introduced by Yasui et al. [36] involves an optimization-based method to handle missing data in gene trees using a mixed integer non-linear programming model. The method is designed to impute missing pairwise distances between leaves in gene trees. It uses a two-stage optimization process, where the first stage involves assigning individuals to hypothetical groups or clades based on the available genetic distances, and the second stage focuses on estimating the missing distances.

Yoshida [37] addresses missing data in phylogenetic trees using tropical geometry. The approach constructs a tropical polytope from known gene trees that are structurally close to the incomplete tree, then projects the incomplete tree onto this polytope in a tropical metric space to estimate the missing parts. The method is restricted to equidistant trees, where all leaves are equidistant from the root. Other approaches address missing data in gene trees using quartet-based optimization or imputation techniques based on tree topology [18,19,22], but do not focus on pairwise tree completion for comparison.

### Our contributions

In this work, we introduce a novel algorithm named *k-Nearest Common Leaves* (*k*-NCL), designed to complete phylogenetic trees that are defined over distinct but overlapping sets of taxa. Our contributions can be summarized as follows. (1) We incorporated branch lengths into the tree completion process, resulting in completed trees that represent both structural and evolutionary relationships, in contrast to topology-only approaches. (2) We introduced a scaling-based strategy to account for differences in evolutionary rates between the two trees. (3) We designed the *k*-NCL algorithm to be independent of any specific distance metric, i.e., it does not assume or optimize for a predefined tree distance, such as RF or geodesic distance in BHV space. (4) We provide a Python implementation of the *k*-NCL algorithm, with open-source code available on GitHub. (5) We evaluated the proposed method on multiple biological datasets (amphibians, birds, mammals, and sharks), where the trees have partial taxon overlap.

## 2 Preliminaries and notation

Let *T* be a rooted phylogenetic tree. Let *V* (*T*) be its set of nodes (both internal and terminal). The leaves (or terminal nodes) of *T*, each denoted by *l*, represent the individual species or taxa under study, and their collection is denoted by *L*(*T*) ⊂ *V* (*T*). The branches (or edges), denoted by *E*(*T*), connect the nodes and represent evolutionary relationships. We assume that each branch *e* = (*u, w*) ∈ *E*(*T*) has an associated strictly positive length between nodes *u* and *w*, indicating the amount of evolutionary change or the time that has passed along that lineage.

### Definition 1

**(Distance between nodes).** *Let u and w be two nodes (internal or terminal) in T*. *The distance between nodes u and w, denoted by d*^(*T*)^(*u, w*), *is defined as the cumulative sum of branch lengths along the unique path connecting u to w*.

### Definition 2

**(Common and distinct leaves).** *In two phylogenetic trees T*_1_ *and T*_2_ *with different but overlapping sets of taxa, the term common leaves refers to the leaves that are present in both trees. The set of common leaves shared by trees T*_1_ *and T*_2_ *is denoted as CL*(*T*_1_, *T*_2_) = *L*(*T*_1_) ∩ *L*(*T*_2_). *Conversely, leaves present in one tree but absent in the other are termed distinct leaves, with the set of such leaves for tree T*_1_ *related to tree T*_2_ *denoted as DL*(*T*_1_|*T*_2_) = *L*(*T*_1_)*\L*(*T*_2_).

Throughout the paper, we assume that |*CL*(*T*_1_, *T*_2_) | ≥ 2 in order for pairwise distances over common leaves to be defined. Let idx(·) denote the rank of an item in a predetermined linear order, defined separately on the common leaves and the original branches of each input tree. (i) Let *CL*(*T*_1_, *T*_2_) be ordered by ascending lexicographic order of taxon labels. For any *l* ∈ *CL*(*T*_1_, *T*_2_), let idx(*l*) denote its position in this order. This rank is used only for deterministic tie-breaking and remains fixed throughout the algorithm. (ii) For each input tree *T*_*i*_, a fixed rank idx(*e*) is assigned to every original branch *e* ∈ *E*(*T*_*i*_) via a single depth-first enumeration established once at the start. If an original branch is split during insertion, all resulting sub-branches inherit the same rank idx(*e*).

### Definition 3

**(Maximal distinct-leaf subtree).** *A subtree S of T*_1_ *is called a maximal distinct-leaf subtree if all leaves of S belong to the set DL*(*T*_1_ | *T*_2_), *and S is not strictly contained in any larger subtree of T*_1_ *whose leaves are also entirely included in DL*(*T*_1_|*T*_2_).

Conceptually, a maximal distinct-leaf subtree *S* is the largest possible subtree in *T*_1_ that contains only leaves not shared with *T*_2_, and cannot be extended without including shared leaves. *S* may consist of one or more distinct leaves, i.e., | *L*(*S*) | ≥ 1. The *root* of *S* in *T*_1_ is defined as the lowest common ancestor (LCA) of all its leaves, denoted by 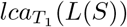. In the special case where *S* contains only a single leaf, its root is that leaf itself. The *root branch* of *S* is the branch that connects the root of *S* to its most immediate ancestor node in the entire tree *T*_1_. We refer to this parent node as the *attachment node* of *S*, denoted 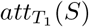. For any given tree *T*, the collection of all its maximal distinct-leaf subtrees is denoted as 𝒮_*T*_.

### Definition 4

**(Nearest common leaves).** *For a distinct-leaf subtree* 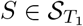, *the set of k nearest common leaves, denoted* 𝒩_*k*_(*S, T*_1_), *consists of the k leaves in T*_1_ *that are also present in T*_2_ *and are closest to the root of S, ordered by increasing distance from it. The parameter k must satisfy* 2 ≤ *k* ≤ |*CL*(*T*_1_, *T*_2_)|.

Let *T* ∈ {*T*_1_, *T*_2_} and let *S* ∈ 𝒮_*T*_. If several common leaves have the same distance as the *k*-th closest leaf to the root of *S*, the ties are resolved using the fixed order of *CL*(*T*_1_, *T*_2_). Specifically, the common leaves *l* ∈ *CL*(*T*_1_, *T*_2_) are ordered by the pair *d*^(*T*)^(*l, lca*_*T*_ (*L*(*S*))), idx(*l*) in lexicographic (increasing) order, and the first *k* leaves are taken. Therefore, |𝒩 _*k*_(*S, T*) | = *k*, and the selection is deterministic.

*Problem 1 (k-NCL tree completion)*. Let *T*_1_ and *T*_2_ be rooted phylogenetic trees with strictly positive branch lengths, leaf sets *L*(*T*_1_) and *L*(*T*_2_), and common leaves *CL*(*T*_1_, *T*_2_) = *L*(*T*_1_)∩*L*(*T*_2_) with |*CL*(*T*_1_, *T*_2_)| ≥ 2. Let *k* ∈ [2, |*CL*(*T*_1_, *T*_2_)|]. The objective is to construct completions 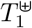 and 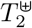 on the unified leaf set *L*(*T*_1_) ∪ *L*(*T*_2_) such that: (i) the restriction of 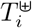 to the leaf set *L*(*T*_*i*_) is identical to *T*_*i*_ in both topology and branch lengths, for *i* ∈ {1, 2}, and (ii) for each *i* ∈ {1, 2} and each maximal distinct-leaf subtree *S* present in *T*_3−*i*_ but absent from *T*_*i*_, *S* is attached to *T*_*i*_ at a location determined from the *k* nearest common leaves of *S* in *T*_3−*i*_. Among all possible attachment locations on the original branches of *T*_*i*_ (including existing internal nodes), the selected location minimizes the discrepancy between distances measured in the target tree and the corresponding position distances induced from *T*_3−*i*_. The completion is obtained by attaching each such subtree in this manner, preserving the original topology and branch lengths on the original leaves.

## 3 Methods

### 3.1 *k*-Nearest Common Leaves algorithm

The *k*-NCL algorithm addresses the tree completion problem by constructing completed trees 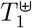 and 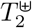 on the unified leaf set *L*(*T*_1_) ∪ *L*(*T*_2_). The method inserts maximal distinct-leaf subtrees from each input tree to the other on the basis of common leaves and adjustment rates, which calibrate branch lengths during subtree insertion.

A high-level description of the *k*-NCL algorithm is given in Appendix D, and Figure 1 presents its main components. The procedure begins with the identification of common leaves, distinct leaves (Definition 2), and maximal distinct-leaf subtrees (Definition 3) in both trees.

**Fig. 1.**
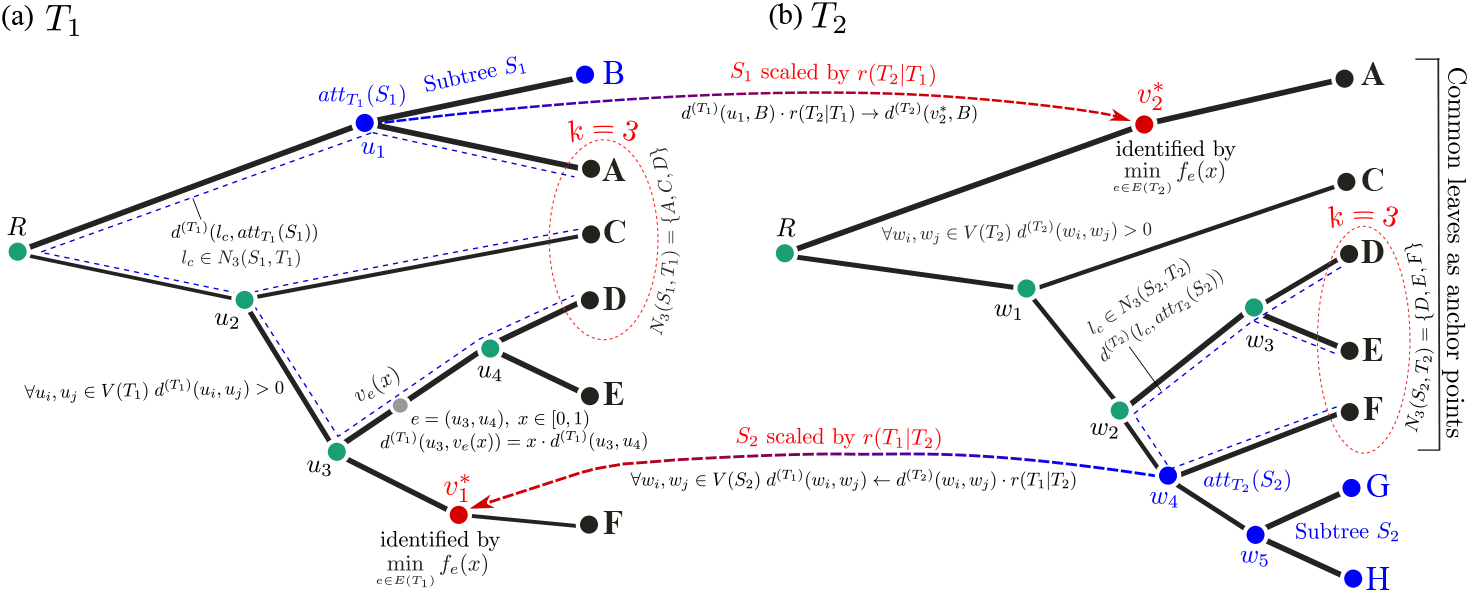
Illustration of the main parts of the *k*-NCL algorithm. The common leaves of trees *T*_1_ and *T*_2_ are marked in bold black. The distinct leaves of each tree are highlighted in blue. Tree *T*_1_ has one one-leaf maximal distinct-leaf subtree {B}. Tree *T*_2_ includes one two-leaf maximal distinct-leaf subtree {G, H}. The value *k* = 3 is chosen. The dashed blue lines represent the position distances from the attachment point of a maximal distinct-leaf subtree to the selected nearest common leaves. The red nodes 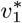 and 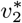 are the insertion points for the corresponding subtrees. The gray candidate node *v*_*e*_(*x*) in *T*_1_ represents an example of the parameterization setting used to find an optimal insertion point along the original branches. Arrows show the addition of maximal distinct-leaf subtrees, with adjusted branch lengths, from one tree to another.

#### Remark 1.

For *i* ∈ {1, 2}, the set 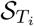 relative to *DL*(*T*_*i*_ | *T*_3−*i*_) can be computed by a post-order traversal of *T*_*i*_. A node is marked if all of its descendant leaves belong to *DL*(*T*_*i*_ *T*_3−*i*_). The roots of maximal distinct-leaf subtrees are precisely the marked nodes whose parent, if any, is unmarked. Each such subtree is then recorded together with its root branch.

#### Remark 2.

Subsequent steps of the *k*-NCL algorithm involve a large number of node-to-node distance computations, specifically on the order of *O*(*n*^2^) in total, where *n* is the total number of taxa across both trees. These include the distances between common leaves, distances from common leaves to subtree roots, and evaluations of candidate insertion points across all branches. To reduce the cost of these operations, we associate each tree with a distance oracle [31], which is rebuilt after every maximal distinct-leaf subtree insertion. The distance oracle is constructed using a depth-first traversal that generates an Euler tour of the tree, along with the depth of each visit and the first occurrence index of every node. A sparse table is built over the depth array to efficiently answer range minimum queries (RMQ), which support constant-time LCA queries using the method of Bender and Farach-Colton [3]. Based on these LCA results, the distance between any two nodes can be computed in *O*(1).

The tree completion procedure continues by adjusting the branch lengths of the maximal distinct-leaf subtrees, including their root branches, using the corresponding branch adjustment rate (Equation 1).

Let *CL*(*T*_1_, *T*_2_) be ordered and indexed. We introduce a global adjustment rate *r*(*T*_1_ | *T*_2_), which is defined as the ratio between the total pairwise distances over all common leaves in *T*_1_ and the corresponding total pairwise distances in *T*_2_ (Equation 1). This rate represents the relative scaling of branch lengths between the two trees based on their shared taxa.

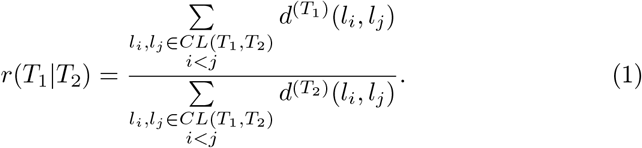

Next, all branch lengths (including the root branch) in the maximal distinct-leaf subtree 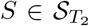, which is to be inserted into *T*_1_, are scaled using the adjustment rate *r*(*T*_1_ | *T*_2_). For each branch (*u, w*) ∈ *E*(*S*) with *E*(*S*) ⊆ *E*(*T*_2_), the adjusted branch length is given by Equation 2.

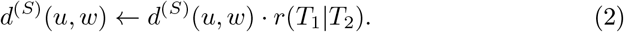

This adjustment is crucial for preserving the phylogenetic relationships and distances within the subtree as it is integrated into the new tree structure. However, the original branch lengths in the initial trees remain unaltered.

Subsequently, the *k* nearest common leaves, 𝒩_*k*_(*S, T*_2_) (Definition 4) are identified for each subtree 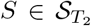 (see Figure 1). Given a selected common leaf, *l*_*c*_ ∈ 𝒩_*k*_(*S, T*_2_), the leaf-based adjustment rate, 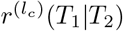, is computed as the ratio of cumulative distances from *l*_*c*_ to all other common leaves in *T*_1_ relative to the corresponding sum in *T*_2_ (Equation 3).

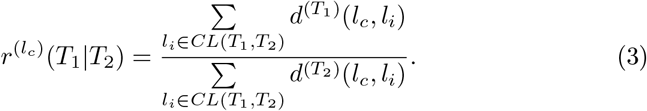

The insertion process is initiated with *T*_1_, and 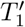 is created as the insertion target for maximal distinct-leaf subtrees from *T*_2_. Subtrees are iteratively inserted into 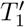 until all subtrees from 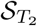 have been added, at which point 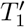 becomes the completed tree 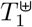. The notation 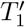 emphasizes that the original tree *T*_1_ remains unchanged throughout the subtree insertion process.

At a high level, the attachment point is chosen by first using the *k* nearest common leaves of the subtree in the source tree to estimate its position relative to the common leaves in the working tree, and then selecting the location on the original branches of the target tree whose distances to these leaves best match the estimated position.

As part of identifying the optimal insertion point for a subtree 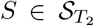, a *position distance* is computed relative to each selected common leaf *l*_*c*_. This distance is calculated as follows (Equation 4).

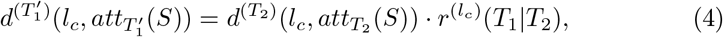

where 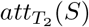 denotes the original attachment node of *S* in *T*_2_, and 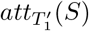 denotes the target attachment location for *S* in 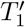.

In order to integrate *S* into 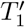, an insertion point *v*^∗^ is selected by minimizing the discrepancy between the observed distances in 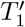 and the computed position distances for the selected common leaves. This procedure is performed as follows.

Every candidate insertion point can lie on an existing node or on a branch connecting two nodes, *u* (the parent node) and *w* (the child node). Fix a candidate branch *e* = (*u, w*) of 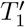. Let *v*_*e*_(*x*) be a point on branch 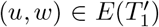 located at a distance 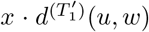 from the parent node *u* along the branch, where *x* ∈ [0, 1) (see Figure 1 (a)). In this parameterization setting, the boundary case *x* = 0 corresponds to the node *u* (an existing node), while *x* approaching 1 corresponds to a point arbitrarily close to the child node *w*. The interval [0, 1) is used to avoid double counting existing nodes.

Then, the *observed distance* from a selected common leaf *l*_*c*_ ∈ 𝒩_*k*_(*S, T*_2_) to the candidate point *v*_*e*_(*x*) is given by Equation 5.

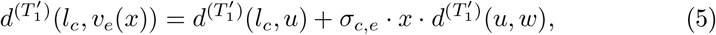

where 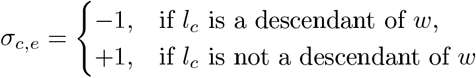.

The discrepancy between the observed distance and the position distance is measured using the following objective function (Equation 6).

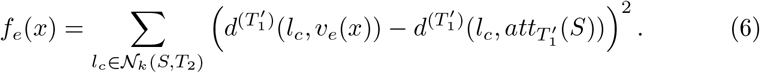

In Equation 6, the sum is taken over *l*_*c*_ ∈ 𝒩_*k*_(*S, T*_2_) (the *k* closest common leaves to *S* in *T*_2_), thus *k* sets the number of terms contributing to *f*_*e*_(*x*) and controls how local versus broader neighborhood information influences the placement. Since *f*_*e*_(*x*) aggregates *k* leaf-wise discrepancies, increasing *k* reduces sensitivity to any single leaf while increasing computation proportionally to *k*.

The function *f*_*e*_(*x*) is quadratic in *x*, and its minimum can be found by taking the derivative with respect to *x*, setting the derivative to zero, and solving for the optimal 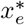. The resulting value 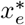 determines the candidate insertion point on the branch as 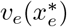.

For each original branch *e* in *T*_1_, as represented in 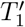, its unique per-branch minimizer 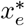 and minimum value 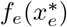 are computed. Let *f*_min_ be the smallest 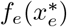 among all original branches. Among all candidates that attain *f*_min_, the insertion point is chosen by minimizing 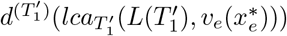. If a tie remains, the candidate on the original branch with the smallest fixed rank idx(*e*) is chosen. This yields a unique insertion point. Then *S* with adjusted branch lengths is placed at this insertion point (see Figure 1).

This process is repeated for each maximal distinct-leaf subtree 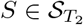 until all have been inserted into 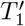, resulting in the final completed tree 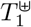. The same procedure is then applied to each subtree from 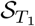 for insertion into a copy of *T*_2_ to construct the completed tree 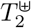. The algorithm outputs two completed phylogenetic trees, 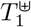 and 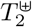, each defined on the union set of taxa of the initial trees 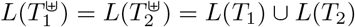. A practical illustration of the algorithm is provided in Section B of the Appendix.

### 3.2 Properties

The following theorem, lemma, and propositions represent several properties of the *k*-NCL tree completion algorithm. Due to the page constraints, the detailed proofs are provided in Section A of the Appendix.

#### Theorem 1

**(Time complexity for fixed *k*).** *Let T*_1_ *and T*_2_ *be phylogenetic trees defined on different but overlapping sets of taxa with L*(*T*_1_) *and L*(*T*_2_), *respectively. For a fixed k* ∈ [2, |*CL*(*T*_1_, *T*_2_)|], *the k-NCL algorithm completes both trees on the combined taxa set L*(*T*_1_)∪*L*(*T*_2_) *in O*(*n*^2^), *where n* = |*L*(*T*_1_)∪*L*(*T*_2_)|.

#### Lemma 1

**(Time complexity for arbitrary *k*).** *For any arbitrary value k* ∈ [2, |*CL*(*T*_1_, *T*_2_)|] *(i*.*e., k is not fixed), the k-NCL algorithm completes both trees on L*(*T*_1_) ∪ *L*(*T*_2_) *in O*(*n*^2^*k*) *time*.

#### Proposition 1

**(Preservation of original distances and topology).** *Let T*_1_ *and T*_2_ *be rooted phylogenetic trees with branch lengths, defined on overlapping leaf sets L*(*T*_1_) *and L*(*T*_2_), *respectively. Let* 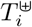 *denote the completed tree produced by the k-NCL algorithm for i* ∈ {1, 2}. *Then, for each i, the completed tree* 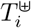 *preserves both the pairwise distances and the topology of the original tree T*_*i*_. *That is, for every pair of original leaves* 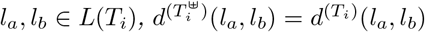. *Furthermore, the completion process preserves the structure of the input tree in the sense that removing the added leaves from* 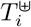 *recovers a tree topologically isomorphic to T*_*i*_.

#### Proposition 2

**(Symmetry in tree completion).** *Let T*_1_ *and T*_2_ *be two phylogenetic trees defined on different but overlapping sets of taxa, and let k* ∈ [2, | *CL*(*T*_1_, *T*_2_) |] *be fixed. The k-NCL tree completion process is symmetric with respect to the input trees. In particular, the algorithm inserts maximal distinct-leaf subtrees from T*_2_ *into T*_1_ *and, symmetrically, maximal distinct-leaf subtrees from T*_1_ *into T*_2_. *Therefore, applying the algorithm to either input order*, (*T*_1_, *T*_2_) *or* (*T*_2_, *T*_1_), *results in the same completed trees* 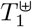 *and* 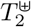, *appearing in the corresponding output order as* 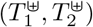 *and* 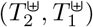.

#### Proposition 3

**(Uniqueness of tree completion in the** *k***-NCL algorithm).** *Let T*_1_ *and T*_2_ *be input phylogenetic trees with strictly positive branch lengths, and let k* ∈ [2, | *CL*(*T*_1_, *T*_2_) |] *be fixed. Then the completed trees* 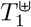 *and* 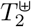 *produced by the k-NCL algorithm are uniquely determined by T*_1_, *T*_2_, *and k, regardless of the order in which maximal distinct-leaf subtrees are inserted*.

#### Proposition 4

**(Applicability to binary and non-binary trees).** *Let T*_1_ *and T*_2_ *be rooted phylogenetic trees with strictly positive branch lengths, defined on overlapping leaf sets L*(*T*_1_) *and L*(*T*_2_), *respectively. Trees T*_1_ *and T*_2_ *may be binary or non-binary (multifurcating). Let k* ∈ [2, | *CL*(*T*_1_, *T*_2_) |]. *Then, the k-NCL tree completion algorithm is well defined for* (*T*_1_, *T*_2_, *k*) *without requiring the trees to be binary. All results in Section 3.2 (polynomial-time complexity, preservation of original distances and topology, symmetry, and uniqueness) remain valid for both binary and non-binary input trees*.

## 4 Results and discussion

### 4.1 Methodology for biological data simulation

The methodology for obtaining biological data of phylogenetic trees with different but overlapping taxa involved several main steps. The biological data used in this evaluation part were sourced from the VertLife website [32], which offers a method for acquiring tree distributions with specific subsets of taxa. The tool prunes a comprehensive dataset to a smaller subset and samples trees from it.

Four different taxonomic groups were selected for analysis, including Amphibians, Birds, Mammals, and Sharks. The number of species in the entire datasets varied across groups, with 7239 species of Amphibians [12], 9993 species of Birds [13], 5911 species of Mammals [32], and 1192 species of Sharks [28]. Each group was first reduced to one representative species per genus. A random sample of *n* species was then drawn from this genus-filtered pool. From these selected species, ten overlapping subsets were generated, with overlap levels ranging from 10% to 90%. The corresponding pruned tree sets were then retrieved from VertLife and combined into the final overlapping datasets.

To introduce diversity into the datasets, different numbers of unique species were selected for each group. In this study, two versions of each biological dataset were prepared (see also Table 2 in the Appendix). Datasets A include 100 phylogenetic trees per group and are used for evaluating the influence of the parameter *k* in the *k*-NCL tree completion algorithm. Datasets B contain a larger number of trees (170 for Amphibians, 100 for Birds, 140 for Mammals, and 100 for Sharks) and are used for the comparative analysis of distance metrics between completed trees. These datasets, along with a more detailed description and the script used to prepare the data, can be found in the project repository on GitHub.

**Table 1.**
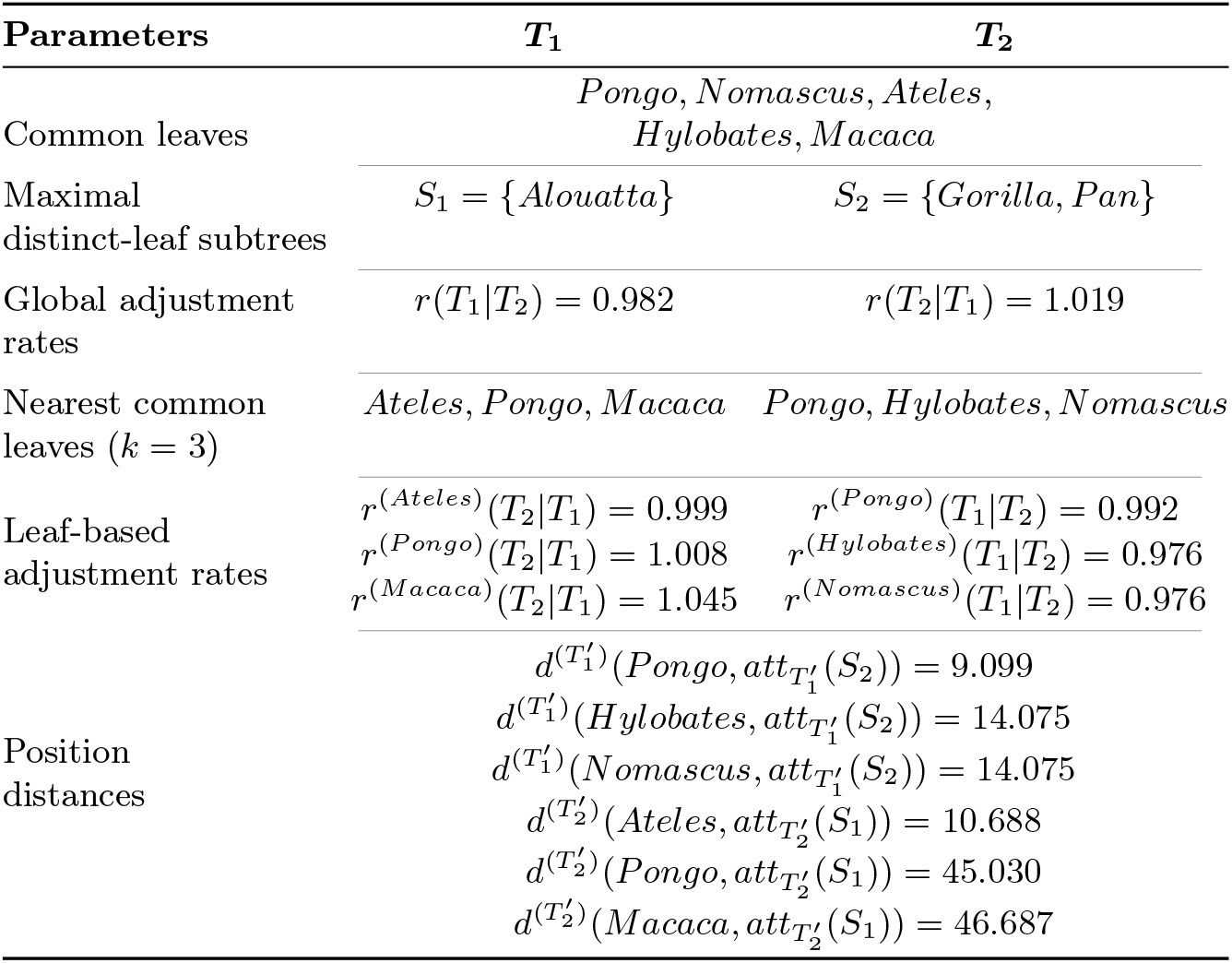
Computed values required for the tree completion process by the *k*-NCL algorithm, with 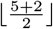 (see the evaluation part results and recommendations). All values are rounded to three decimal places.

**Table 2.** Summary of the four biological datasets used in this study: Amphibians, Birds, Mammals, and Sharks. Each dataset consists of a set of individual phylogenetic trees (e.g., 100 trees for each taxonomic group in Datasets A). For evaluation, all unique tree pairs are considered from each dataset, resulting in 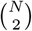 comparisons per group, where *N* is the number of trees.

| Groups | Number of trees |  | Unique species | Source |
| --- | --- | --- | --- | --- |
|  | Datasets A | Datasets B |  |  |
| Amphibians | 100 | 170 | 120 | [12] |
| Birds | 100 | 100 | 135 | [13] |
| Mammals | 100 | 140 | 105 | [32] |
| Sharks | 100 | 100 | 95 | [28] |

### 4.2 Evaluation setup

In this part of the study, four distance metrics are applied, involving topology-only and topology and branch length comparisons, along with pruning and completion-based approaches. The Branch Score Distance (BSD) [14] is used in two variants in this section. BSD(*k*-NCL) measures the distance between trees completed using the *k*-NCL algorithm, while BSD(-) computes the distance after pruning both trees to their common taxa. Two versions of the RF distance [25] are utilized, each based on a different tree completion strategy. The first, RF(+), is computed on trees completed using the method proposed in [35]. This metric only considers topological differences and ignores branch lengths, making it suitable for evaluating tree structure without regard to evolutionary distances. The second version, RF(*k*-NCL), applies the classical RF distance to trees completed using the *k*-NCL method. While the RF metric itself remains purely topological, this variant reflects how the *k*-NCL algorithm affects tree topology.

For this analysis, tree pairs are categorized by their levels of overlap, which are quantified using the Jaccard coefficient [23]. For the results in this section, overlap levels are binned into intervals of width 0.1, with each level (e.g., 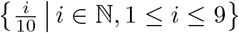 representing the center of a bin. For instance, the level 0.1 corresponds to the interval [0.05, 0.15), 0.2 to [0.15, 0.25), continuing up to 0.9, which corresponds to [0.85, 0.95).

### 4.3 Effect of the parameter *k*

Let *N*_*cl*_ denote the number of common leaves, i.e., *N*_*cl*_ = | *CL*(*T*_1_, *T*_2_) |. For each tree pair, the BSD(*k*-NCL) distance is computed for a range of *k* values. These include edge cases such as *k* ∈ [2, 3, *N*_*cl*_ − 1, *N*_*cl*_], along with intermediate values, including 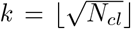 and 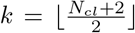. Tree pairs are grouped into intervals based on overlap level, and for each group, the average BSD(*k*-NCL) is plotted as a function of *k*. This reveals how the distance changes with varying *k* values. These results are illustrated in Figure 2.

**Fig. 2.**
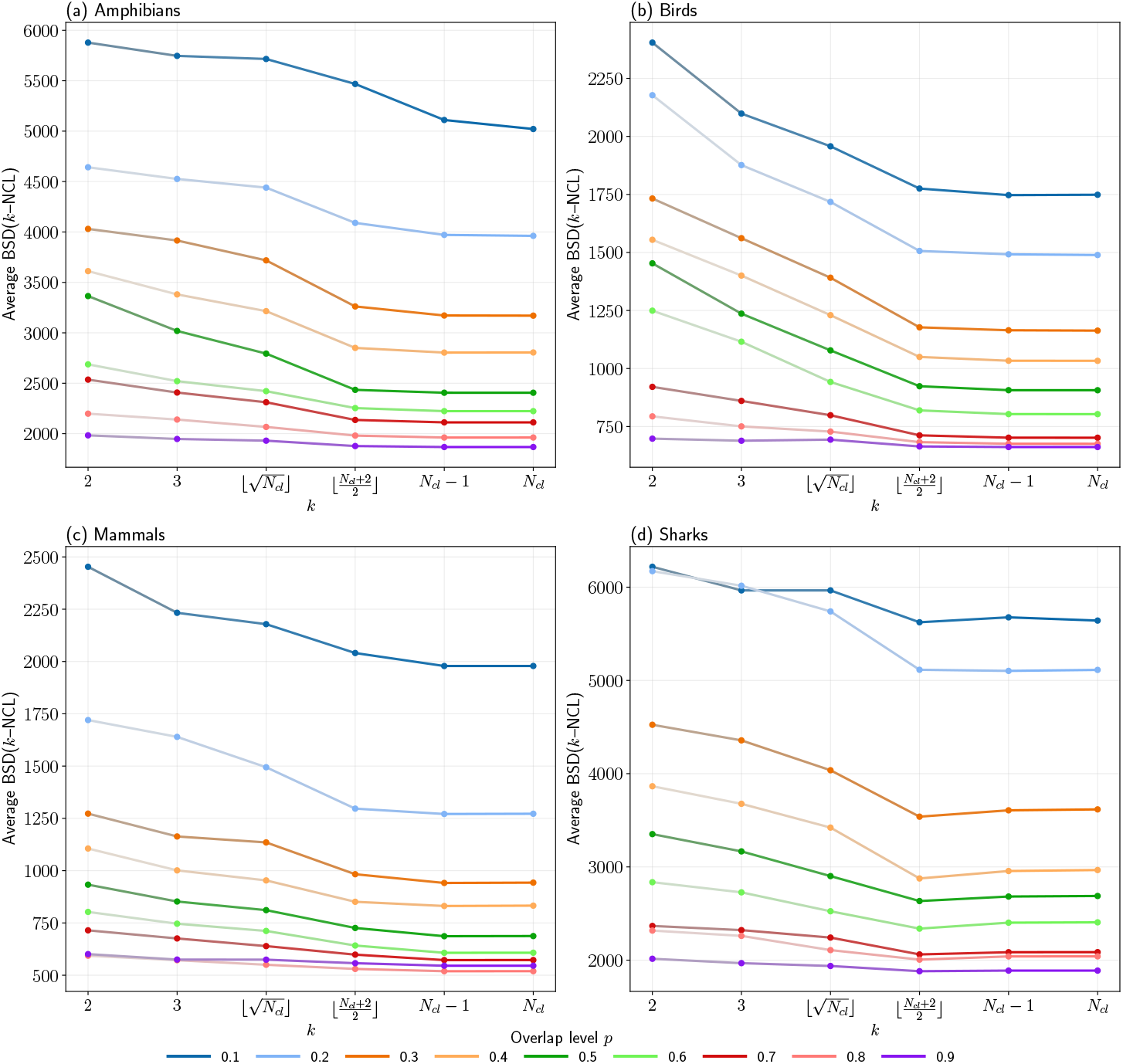
Average BSD(*k*-NCL) distance versus *k* values for various levels of overlap (Datasets A). Edge cases (*k* = [2, 3, *N*_*cl*_ − 1, *N*_*cl*_]) and intermediate cases (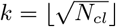 and 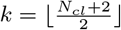) are used to complete tree pairs using the *k*-NCL algorithm.

The results show a consistent and decreasing trend in general. Across all four datasets, the average BSD(*k*-NCL) distance decreases as *k* increases. The decrease is gradual at first and then largely flattens as *k* approaches 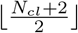 and *N*_*cl*_. This pattern holds across different overlap levels, with one exception for Sharks, where the lowest average BSD(*k*-NCL) occurs at 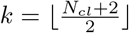 rather than at the largest *k*.

Analysis of the optimal *k* for individual tree pairs supports this trend. Empirically, 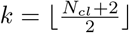 can be used as a suitable default. It achieves nearly minimal distances, and larger *k* provides only minor additional improvements, on average. Accordingly, 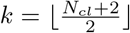 is used as the default value in subsequent evaluations in this study.

### 4.4 Comparison between completion and pruning

In order to evaluate the agreement between BSD(*k*-NCL) and BSD(-), three scenarios are defined based on how each distance interprets the similarity of two trees, *T*_1_ and *T*_2_, relative to a reference tree *T* ^∗^. A supertree with branch lengths was constructed as a reference tree for this evaluation part. The overall approach to obtaining a reference tree for each dataset involved first inferring the supertree topology, followed by assigning branch lengths by averaging values across consistent splits from the source trees. The spectral cluster supertree algorithm [20] was applied to build the supertree from a collection of input trees, each defined on different but overlapping subsets of taxa and containing branch length information. To assign branch lengths to the supertree, a post-processing step was performed. Each internal branch (split) of the inferred supertree was matched against the corresponding splits in the original input trees based on their bipartition structure. For every matching split identified, the associated branch lengths were recorded. The final branch length for each supertree branch was calculated as the arithmetic mean of all matching branch lengths across the source trees.

The following three scenarios are similar to the evaluation procedures described in [35]. Scenario 1 includes cases where the two metrics provide opposite conclusions. One metric indicates that *T*_1_ is more similar to the reference tree *T* ^∗^, while the other indicates that *T*_2_ is more similar to *T* ^∗^. Scenario 2 identifies cases where BSD(-) gives equal distances for both trees, but BSD(*k*-NCL) assigns different distances. This indicates that BSD(*k*-NCL) detects a difference between the trees that BSD(-) does not. Scenario 3 captures the opposite of Scenario 2, where BSD(*k*-NCL) gives equal distances, but BSD(-) differentiates the two trees. These scenarios are used to identify conflicting pairs of trees under each comparison, depending on their level of leaf overlap. Results comparing BSD(*k*-NCL) and BSD(-) are shown in Figure 3.

**Fig. 3.**
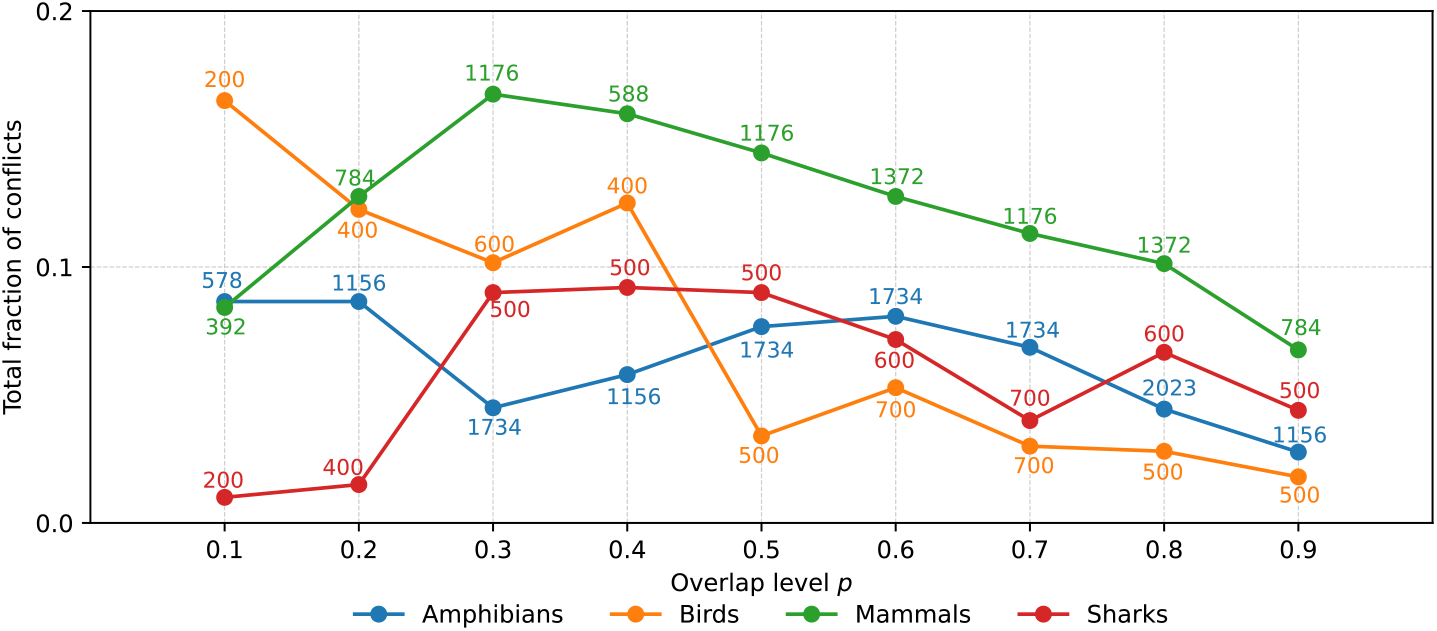
Fraction of conflicting tree pairs for BSD(*k*-NCL) versus BSD(-) for all scenarios (Datasets B). 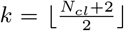 is used for BSD(*k*-NCL). Numbers above points show the number of tree pairs in each bin.

Across all four datasets, 8.02% of tree pair comparisons revealed conflicts between BSD(*k*-NCL) (completion) and BSD(-) (pruning). Among those conflicts, 98.9% are Scenario 1, while Scenarios 2 and 3 contribute only 0.69% and 0.41% of conflicts, respectively. The conflict rates by dataset are as follows: 12.4% for Mammals, 6.2% for Amphibians, 6.5% for Birds, and 6.2% for Sharks. Scenario 1 dominates in each dataset (99.5% for Amphibians, 97.3% for Birds, 98.6% for Mammals, and 100% for Sharks).

Conflict levels vary with leaf overlap *p*. Aggregating across datasets, the conflict fraction peaks around *p* = 0.4 (9.7%), and falls to 3.9% by *p* = 0.9 from 8.6% at *p* = 0.1. Dataset specific patterns align with Figure 3. Birds decline steadily from 16.5% at *p* = 0.1 to 1.8% at *p* = 0.9. Mammals show a peak of 16.8% at *p* = 0.3. Amphibians are generally lower, decreasing from 8.7% at *p* = 0.1 to 2.8% at *p* = 0.9. Sharks show a moderate peak at *p* = 0.4 (9.2%).

The median total conflict fraction is highest at *p* = 0.4 (the median is 10.8%) and decreases to 3.6% at *p* = 0.9, indicating both a reduction in conflict and convergence among datasets as overlap increases. It can be concluded that conflicts between completion-based BSD(*k*-NCL) and pruning-based BSD(-) are infrequent overall (≈ 8%) and occur predominantly as Scenario 1. The risk of conflict is highest at low and medium levels of overlap (*p* ≤ 0.4), notably for Mammals, and decreases with increasing overlap (the median across datasets is approximately 3% − 4% at *p* = 0.9). In practice, pruning can discard information when the overlap is limited. Therefore, BSD(*k*-NCL) may provide a more informative comparison at low and medium levels of overlap, whereas at high overlap (*p* ≥ 0.8) the two metrics behave similarly.

### 4.5 Tree completion methods comparison

In order to evaluate the proposed *k*-NCL tree completion method in comparison to the RF(+) tree completion approach, a clustering-based analysis was conducted. RF(+) was selected as the main external baseline because, to the best of our knowledge, it is the only previously published method outside our earlier method in [14] that directly addresses the tree completion problem, namely the construction of completed trees on the union of taxa from two overlapping input trees. The *k*-NCL method extends the approach introduced in [14]. Other related approaches either define distances that require identical taxon sets, or operate via pruning or implicit embeddings without explicitly returning completed trees. For this reason, RF(+) is the only direct external algorithmic baseline for tree completion considered here. It is important to note that both RF(+) and *k*-NCL attach maximal distinct-leaf subtrees, and the primary difference is the placement criterion. RF(+) is topology-driven, whereas *k*-NCL selects attachment points by minimizing a branch-length-based least-squares discrepancy. This comparison is designed to highlight the differences between topology-based and branch length-based tree completion outputs. The evaluation aims to quantify the ability of three tree distance metrics (RF(+), RF(*k*-NCL), and BSD(*k*-NCL)) to identify intended clusters from sets of partially overlapping phylogenetic trees of Amphibians, Birds, Mammals, and Sharks.

For each species group, five empirical base trees were selected, each defining one reference cluster. The base trees were selected such that the leaf set overlap across the resulting clusters ranged from 10% to 90%. From each base tree, four perturbed variants were simulated with AsymmeTree [27], resulting in five related trees per cluster and 25 trees per species group in total. This setup was designed as a controlled benchmark with known cluster membership, rather than as a simulation of fully independent empirical trees. Full simulation details and parameter settings are provided in the project repository.

For every unique tree pair within each cluster dataset, RF(+) and *k*-NCL completion methods are applied. RF distances are calculated for the trees completed with RF(+), and both RF(*k*-NCL) and BSD(*k*-NCL) distances are computed for trees completed with *k*-NCL 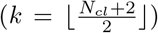. The resulting pairwise distance matrices for each of the three metrics (RF(+), RF(*k*-NCL), and BSD(*k*-NCL)) are visualized in Figure 4. Heatmaps and boxplots illustrate how well each tree completion method reveals the expected clustering structure within the datasets.

**Fig. 4.**
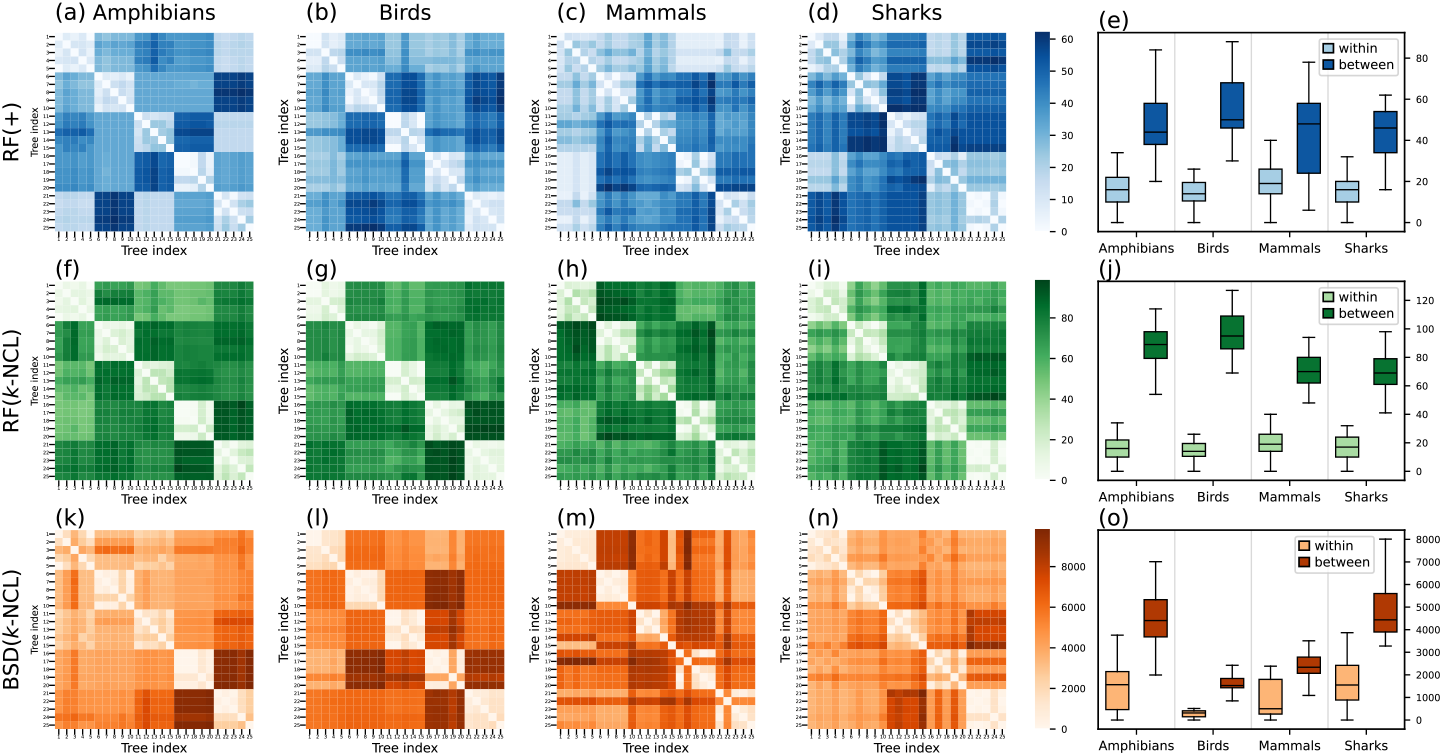
Pairwise distance heatmaps and summary boxplots for four species groups and three tree distance metrics. Each row corresponds to a distance metric, including RF(+), RF(*k*-NCL), and BSD(*k*-NCL). The first four columns correspond to a species group of Amphibians, Birds, Mammals, and Sharks. There are 25 trees per group, arranged into five clusters of five trees. A value of 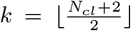 was used for tree completion in RF(*k*-NCL) and BSD(*k*-NCL).

In the RF(+) row panels (a) to (d) of Figure 4, the blocks are present, but they are less distinct. Panel (c), which represents Mammals, shows fuzzy boundaries with visible overlap between clusters. The RF(+) boxplots (e) also reveal overlap. In the RF(*k*-NCL) row panels (f) to (i), the block diagonals are clearly defined, with very light within-cluster cells and darker between-cluster bands. The corresponding boxplots (j) demonstrate a distinct gap. In the BSD(*k*-NCL) panels (k) to (n), the blocks are visible, however the edges are less distinct than in the RF (*k*-NCL) panels. The boxplots in panel (o) show a small but noticeable overlap.

Cluster separation was quantified from the 25×25 pairwise distance matrices for each method and species group, using the known clustering structure of five clusters, each consisting of five trees. The distributions of within-cluster and between-cluster distances were extracted for each method and species group. The silhouette method [26] and the Dunn index [9] were utilized to evaluate cluster separation. The silhouette coefficient for a single tree is computed from a distance matrix by defining *a* as the mean distance of that tree to all other trees within its own cluster, and *b* as the mean distance to all trees in the nearest other cluster. The coefficient is then given by (*b* − *a*)*/* max(*a, b*). The overall silhouette score is the average of these coefficients across all trees. The Dunn index equals the minimum inter-cluster distance divided by the maximum intra-cluster diameter. Larger values indicate better cluster separation for all metrics. These values are presented in Table 3 in the Appendix.

**Table 3.**
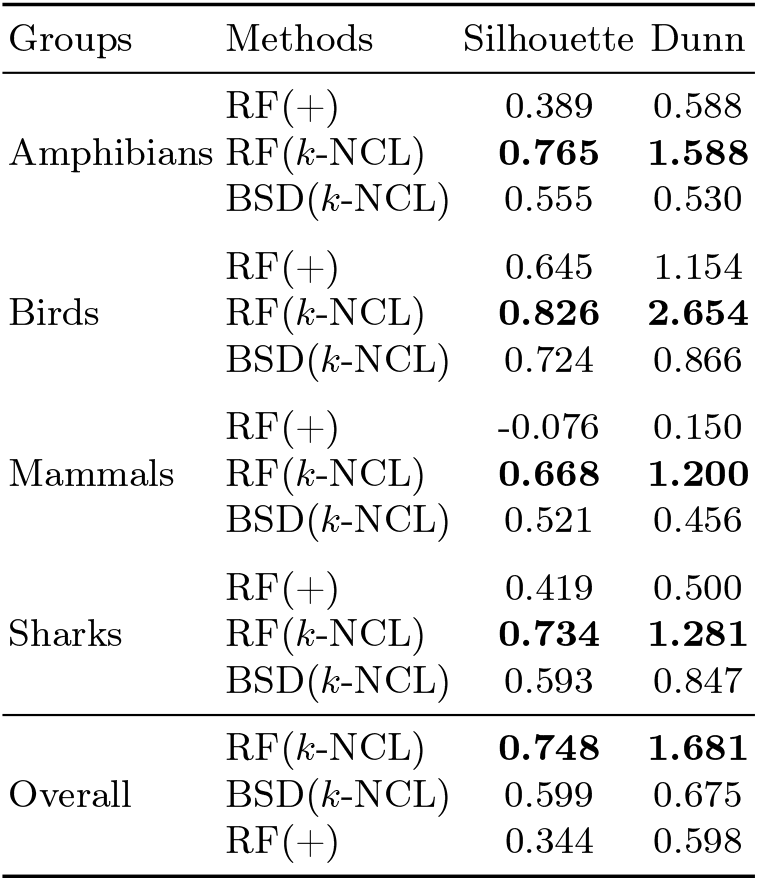
Cluster separation metrics (silhouette and Dunn). The highest values in each species group and for each metric are highlighted in bold. The final *Overall* block ranks methods by their aggregated values across all groups. Values are rounded to three decimals.

RF(*k*-NCL) shows perfect ordering of distances in all species groups and the largest silhouette scores and Dunn indices (Amphibians silhouette 0.765 and Dunn 1.588, Birds silhouette 0.826 and Dunn 2.654, Mammals silhouette 0.668 and Dunn 1.200, Sharks silhouette 0.734 and Dunn 1.281). The boxplots (Figure 4) display a gap between the light within-cluster boxes and the dark between-cluster boxes under RF(*k*-NCL) with no overlap. BSD(*k*-NCL) provides clear separation that is close to RF(*k*-NCL) in Birds and Sharks, but with moderate overlap and lower silhouettes and Dunn indices (Birds silhouette 0.724 versus 0.826 and Dunn 0.866 versus 2.654, Sharks silhouette 0.593 versus 0.734 and Dunn 0.847 versus 1.281), while still outperforming RF(+) in Mammals (silhouette 0.521 versus −0.076 and Dunn 0.456 versus 0.150). RF(+) shows good results for Birds, but demonstrates noticeable overlap in other groups and weaker separation of Mammal clusters as indicated by a negative silhouette and the smallest Dunn value (Birds silhouette 0.645 and Dunn 1.154, Mammals silhouette −0.076 and Dunn 0.150). The most robust method for cluster identification and separation in these data is RF(*k*-NCL) overall and within Amphibians, Birds, Mammals, and Sharks (overall silhouette 0.748 and Dunn 1.681). BSD(*k*-NCL) is a second choice, and RF(+) ranks third (overall silhouette 0.599 and Dunn 0.675 for BSD(*k*-NCL) and overall silhouette 0.344 and Dunn 0.598 for RF(+)).

These results suggest that the *k*-NCL method, by integrating both topology and branch lengths, can enhance cluster separation for overlapping trees. Furthermore, both topology-based (RF) and branch length-based (BSD) distances benefit from the tree completion results provided by the *k*-NCL method.

## 5 Conclusion

This study presents the *k*-Nearest Common Leaves (*k*-NCL) algorithm, designed for completing phylogenetic tree pairs that are defined on different but overlapping taxa. The method uses branch lengths and topology to insert non-common leaves while preserving evolutionary distances and topological structure. The resulting completions extend both trees to the same taxon set while retaining their original structure. The algorithm runs in *O*(*n*^2^) time in the size of the union of leaf sets (Theorem 1) and satisfies the preservation of original distances and topology (Proposition 1), symmetry (Proposition 2), and uniqueness (Proposition 3). It applies to binary and non-binary trees (Proposition 4). The proposed algorithm is evaluated on datasets of amphibians, birds, mammals, and sharks (Section 4.1), and improves phylogenetic tree clustering compared to RF(+) completions (Section 4.5).

Future work will include additional analysis of how the choice of the parameter *k* affects the quality of tree completion by relating it to a purely topological distance metric and studying its impact on the robustness and stability of the tree completions. On the algorithmic side, further optimizations of the algorithm to improve scalability, as well as extensions to completing collections of more than two trees, are promising directions. In particular, the approach can be extended to a set of partially overlapping phylogenetic trees in order to obtain a completed set, where each tree is defined on a combined set of taxa. Finally, additional evaluation experiments on larger and more diverse datasets will help to better characterize the behavior of the *k*-NCL method across a wide range of evolutionary scenarios.

## Appendix

### A Proofs for Section 3.2 (Properties)

The following theorem, lemma, and propositions represent several properties of the *k*-NCL tree completion algorithm.

#### Theorem 1

**(Time complexity for fixed** *k*). *Let T*_1_ *and T*_2_ *be phylogenetic trees defined on different but overlapping sets of taxa with L*(*T*_1_) *and L*(*T*_2_), *respectively. For a fixed k* ∈ [2, |*CL*(*T*_1_, *T*_2_)|], *the k-NCL algorithm completes both trees on the combined taxa set L*(*T*_1_)∪*L*(*T*_2_) *in O*(*n*^2^), *where n* = |*L*(*T*_1_)∪*L*(*T*_2_)|.

*Proof*. Let *n* = | *L*(*T*_1_) ∪ *L*(*T*_2_) | represent the total number of taxa in the combined leaf set of *T*_1_ and *T*_2_. We assume that *k* is a fixed constant, independent of *n*. Since each phylogenetic tree *T*_*i*_ (for *i* ∈ {1, 2}) is rooted, the number of internal nodes in each tree is at most | *L*(*T*_*i*_) | − 1. Thus, each *T*_*i*_ has at most 2 | *L*(*T*_*i*_) | − 1 nodes (internal and terminal) and 2 | *L*(*T*_*i*_) | − 2 branches. As | *L*(*T*_*i*_) | ≤ *n*, a traversal of *T*_*i*_ can be performed in *O*(*n*).

At the beginning of the tree completion process, the set of common leaves, *CL*(*T*_1_, *T*_2_) = *L*(*T*_1_) ∩ *L*(*T*_2_), and the set of distinct leaves, *DL*(*T*_3−*i*_|*T*_*i*_) = *L*(*T*_3−*i*_) \ *CL*(*T*_1_, *T*_2_), are computed. Since the total number of taxa is *n* = | *L*(*T*_1_) ∪ *L*(*T*_2_) |, each of these sets contains at most *n* elements. To compute the intersection and set difference efficiently, all leaf labels from one set (e.g., *L*(*T*_1_)) are inserted into a hash set, which supports expected constant-time lookups. The other leaf set (e.g., *L*(*T*_2_)) is then iterated to check which elements are present or absent in the hash set. Constructing the hash set, performing these lookups, and computing the resulting sets of common and distinct leaves each take at most *O*(*n*) time. Thus, the total cost of this step is *O*(*n*).

In order to find maximal distinct-leaf subtrees (Remark 1), the tree is traversed using post-order traversal, visiting each node exactly once. At each node, checks are performed, and the node may be marked or added to the result set, all in constant time. After traversal, the nodes are processed again to collect the roots of valid subtrees, which is done in linear time. Since each step involves only constant work per node, the overall time complexity of this step is *O*(*n*).

Efficient distance queries are achieved by constructing a distance oracle for each working tree (Remark 2). The oracle is based on an Euler tour representation of the tree, which takes *O*(*n*) time to compute via a depth-first traversal. The tour records each visit to a node and tracks its depth (where the depth of a node is defined as the total branch length from the tree root to that node), along with the first occurrence of every node. The Euler tour has linear length and can be constructed in *O*(*n*) time. Constant-time retrieval of LCA between any two nodes is achieved by constructing an RMQ structure over the depth array of the tour. This can be performed in *O*(*n*) time using the algorithm of Bender and Farach-Colton [3]. Therefore, using the precomputed LCA data structure (i.e., the Euler-tour array, its depth array, first-occurrence array, and RMQ table), the distance between any two nodes *u* and *w* can be retrieved in constant time by computing the sum of their depths minus twice the depth of their lowest common ancestor. Thus, the complete distance oracle is built in linear time and supports constant-time distance queries between any pair of nodes.

Each time a subtree is inserted into the working copy, the tree topology changes. The distance oracle is rebuilt after every insertion to account for these changes. Since there are at most *n* such insertions, the cumulative cost of updating all distance oracles across the algorithm is *O*(*n*^2^).

The global adjustment rate *r*(*T*_*i*_ | *T*_3−*i*_) is calculated by comparing the cumulative pairwise distances between all common leaves (Equation 1). Since there are at most *n* common leaves, this computation involves *O*(*n*^2^) distance queries, each answered in constant time via the distance oracle. The total cost of computing this rate is therefore *O*(*n*^2^).

For each maximal distinct-leaf subtree *S*, up to *k* (constant) leaf-based adjustment rates and position distances are computed, as described in Equations 3 and 4. Each such computation involves a linear number of distance queries across the common leaf set. Consequently, each subtree contributes *O*(*n*) to the total runtime, and across all subtrees, the accumulated cost remains *O*(*n*^2^).

For each candidate branch in the target tree, the objective function *f*_*e*_(*x*) (Equation 6) is evaluated. Since the number of such leaves is constant (*k*) and each distance can be retrieved in constant time via the distance oracle, the cost of evaluating *f*_*e*_(*x*) at a candidate point is *O*(1). The minimum of this quadratic function can then be found in constant time. As the number of candidate branches is linear in the tree size, finding the optimal insertion point for one subtree takes *O*(*n*) time. The tie-breaking comparisons are *O*(1) per branch. Summed over all insertions, this step contributes *O*(*n*^2^) to the total running time.

In addition, operations such as scaling branch lengths, tagging nodes, and splitting branches are each linear in the size of the subtree being inserted.

Combining all components, the total runtime of the algorithm is *O*(*n*^2^) for fixed *k*. Therefore, the *k*-NCL algorithm runs in quadratic time.

Theorem 1 states the complexity in the fixed-*k* setting, and the following Lemma 1 gives the corresponding bound when *k* is not assumed to be fixed.

#### Lemma 1

*Proof*. All structural preprocessing steps that do not depend on *k* remain as in Theorem 1. Given that each 𝒩_*k*_(*S, T*_*i*_) has exactly *k* leaves, the selection of *k* nearest common leaves for 𝒩 _*k*_(*S, T*_*i*_) requires scanning the | *CL*(*T*_1_, *T*_2_)| ≤ *n* common leaves and extracting the *k* smallest distances using constant-time distance queries. This step costs *O*(*n*) per subtree and is dominated by the terms below.

The subsequent dependence of the algorithm on *k* occurs in two places. First, up to *k* leaf-based adjustment rates and position distances are computed for each maximal distinct-leaf subtree *S* (Equations 3 and 4). Each such computation performs a linear number of distance queries over the set of common leaves, i.e., *O*(*n*). One subtree contributes *O*(*nk*), and it results in *O*(*n*^2^*k*) for at most *n* insertions.

Second, the objective function *f*_*e*_(*x*) (Equation 6) aggregates discrepancies across the *k* nearest common leaves for each candidate branch during the placement of *S*. A single evaluation of *f*_*e*_(*x*) costs *O*(*k*) with constant-time distance queries from the precomputed distance oracle. Since the number of candidate branches is linear in the tree size, optimizing one insertion takes *O*(*nk*) time, and it is *O*(*n*^2^*k*) for up to *n* insertions.

All remaining operations are dominated by *O*(*n*^2^*k*). Therefore, the total running time of the *k*-NCL algorithm, under the assumption that *k* is not fixed, is *O*(*n*^2^*k*).

#### Proposition 1

**(Preservation of original distances and topology).** *Let T*_1_ *and T*_2_ *be rooted phylogenetic trees with branch lengths, defined on overlapping leaf sets L*(*T*_1_) *and L*(*T*_2_), *respectively. Let* 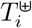 *denote the completed tree produced by the k-NCL algorithm for i* ∈ {1, 2}. *Then, for each i, the completed tree* 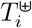 *preserves both the pairwise distances and the topology of the original tree T*_*i*_. *That is, for every pair of original leaves* 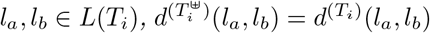. *Furthermore, the completion process preserves the structure of the input tree in the sense that removing the added leaves from* 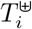 *recovers a tree topologically isomorphic to T*_*i*_.

*Proof*. The *k*-NCL algorithm constructs 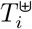 by copying the original tree *T*_*i*_ to obtain 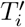, which is used as the basis for inserting maximal distinct-leaf subtrees derived from the other tree. During this process, no changes are made to the existing topology or branch lengths of *T*_*i*_. If a subtree is inserted at a location along a branch (*u, w*), the insertion may involve splitting the branch at a point *v*^∗^, producing two sub-branches. The total length of the original branch is maintained through this operation, as described by Equation 7.

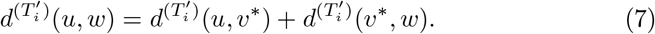

Since the cumulative path length across any affected branch remains unchanged and no other modifications are made to existing paths or nodes in 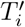 it follows that the distances between any pair of original leaves in *L*(*T*_*i*_) remain the same in 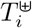, as shown in Equation 8.

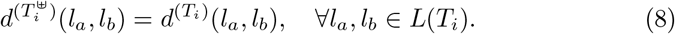

Topologically, the only modifications to the tree involve attaching new subtrees at points along existing branches without altering the branching order or connectivity among the original nodes. Therefore, the subtree induced by *L*(*T*_*i*_) in 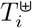 is structurally identical to the original tree *T*_*i*_, meaning that it is topologically isomorphic. Thus, both the pairwise evolutionary distances and the tree structure of the original taxa are preserved within the completed tree, supporting the consistency of the phylogenetic signal.

#### Proposition 2

**(Symmetry in tree completion).** *Let T*_1_ *and T*_2_ *be two phylogenetic trees defined on different but overlapping sets of taxa, and let k* ∈ [2, | *CL*(*T*_1_, *T*_2_) |] *be fixed. The k-NCL tree completion process is symmetric with respect to the input trees. In particular, the algorithm inserts maximal distinct-leaf subtrees from T*_2_ *into T*_1_ *and, symmetrically, maximal distinct-leaf subtrees from T*_1_ *into T*_2_. *Therefore, applying the algorithm to either input order*, (*T*_1_, *T*_2_) *or* (*T*_2_, *T*_1_), *results in the same completed trees* 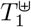 *and* 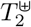, *appearing in the corresponding output order as* 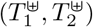 *and* 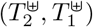.

*Proof*. Let *T*_1_ and *T*_2_ be two phylogenetic trees defined on overlapping but non-identical sets of taxa, and assume the number of common leaves | *CL*(*T*_1_, *T*_2_) | ≥ 2. We analyze each step of the *k*-NCL algorithm and demonstrate its symmetry under the exchange of *T*_1_ and *T*_2_. Let *k* ∈ [2, | *CL*(*T*_1_, *T*_2_) |] be fixed.

The first step involves computing the set of common leaves *CL*(*T*_1_, *T*_2_) and the distinct leaf sets *DL*(*T*_1_ | *T*_2_) and *DL*(*T*_2_ | *T*_1_). These operations are symmetric by definition.

Next, the sets of maximal distinct-leaf subtrees 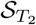 and 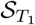 are constructed. Since the procedure for identifying such subtrees depends only on the structure of the input tree and the distinct leaf set, which is itself symmetric under tree exchange.

The global adjustment rate *r*(*T*_1_ | *T*_2_) is computed as defined in Equation 1, and *r*(*T*_2_ | *T*_1_) is its reciprocal. Thus, both rates are derived from the same symmetric structure of pairwise distances between common leaves. Likewise, the subtree scaling in Equation 2 applies the adjustment rate uniformly to the internal branches of the subtrees, and this operation is symmetric in form under tree reversal, differing only in which tree provides the scaling reference.

The selection of the nearest common leaves 𝒩_*k*_(*S, T*_2_) and 𝒩_*k*_(*S*^*′*^, *T*_1_) follows Definition 4, which is purely distance-based and invariant in structure under the reversal of input trees. The computation of leaf-based adjustment rates (Equation 3) is structurally identical in both directions, relying solely on distances among common leaves.

The computation of position distances in Equation 4, along with the observed distances and the construction of the objective function in Equation 6, are all expressed in the same mathematical form. These components depend only on the current target tree, the subtree to be inserted, and the distances among common leaves. Hence, it is structurally symmetric.

Finally, the minimization of the objective function over insertion points on each branch is performed identically in both directions. Since the optimization problem defined by Equation 6 is convex and quadratic in *x*, its analytical solution follows the same procedure for *T*_1_ and *T*_2_.

Overall, each computational and structural step of the algorithm applied to complete *T*_1_ using *T*_2_ is equivalently reproduced in the process of completing *T*_2_ using *T*_1_. Therefore, the tree completion process defined by the *k*-NCL algorithm is symmetric with respect to the input trees.

#### Proposition 3

**(Uniqueness of tree completion in the** *k***-NCL algorithm).** *Let T*_1_ *and T*_2_ *be input phylogenetic trees with strictly positive branch lengths, and let k* ∈ [2, | *CL*(*T*_1_, *T*_2_) |] *be fixed. Then the completed trees* 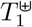 *and* 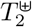 *produced by the k-NCL algorithm are uniquely determined by T*_1_, *T*_2_, *and k, regardless of the order in which maximal distinct-leaf subtrees are inserted*.

*Proof*. For each *i* ∈ {1, 2}, let 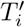 be the working copy of the input phylogenetic tree *T*_*i*_ used by the algorithm to insert maximal distinct-leaf subtrees from *T*_3−*i*_ onto the original branches *e* of 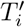 (i.e., branches corresponding to *E*(*T*_*i*_)). For each such original branch, the objective function *f*_*e*_(*x*) (Equation 6) is a strictly convex quadratic function in *x*. Therefore, it has a unique minimizer 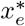. Ties can only occur across different branches if several branches attain the same minimum value.

The established tie-breaking rule first restricts to candidates with the global minimum value of the objective function, then minimizes the distance to the root 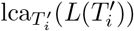, and finally uses the fixed rank idx(*e*) of the underlying original branch. This induces a total order on candidates and yields a unique insertion point.

Since candidates are placed only on the original branches and the distances between the original nodes are preserved (Proposition 1), the candidate set is unaffected by the order of previous insertions. Therefore, the outcome is independent of the subtree insertion order.

Thus, the insertion points for all subtrees and, consequently, the completed trees are uniquely determined by *T*_1_, *T*_2_, and *k*.

#### Proposition 4

*Proof*. For each input tree *T*_*i*_ with *i* ∈ {1, 2}, the distance 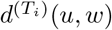 is the path length along the unique path between *u* and *w*, which is well defined in any tree with strictly positive branch lengths. The sets *CL*(*T*_*i*_, *T*_3−*i*_) and *DL*(*T*_*i*_ | *T*_3−*i*_) depend only on leaf labels. The LCA of a non-empty set of nodes exists and is unique in any rooted tree. The definition of a maximal distinct leaf subtree depends only on the ancestor relation and on membership in *DL*(*T*_*i*_ | *T*_3−*i*_). The *k* nearest common leaves are chosen using node to node distances and the fixed index order. None of these constructions require internal nodes to have degree two.

For complexity, even if the internal nodes of a tree are multifurcating, every traversal, Euler tour, and distance oracle construction remains *O*(*n*) in the number of leaves, up to a constant factor. Hence, Theorem 1 and the subsequent Lemma 1 give the same bounds *O*(*n*^2^) for fixed *k* and *O*(*n*^2^*k*) for arbitrary *k* in the non-binary case.

The proofs of Propositions 1, 2, and 3 use only the preservation of edge lengths on original branches, the tree metric property, the existence of LCA, the strict convexity of *f*_*e*_(*x*), and the deterministic tie breaking rule. None of these parts depend solely on binary branching. Hence, *k*-NCL can be applied without modifications to binary and non-binary trees, and all stated properties hold in this general setting.

### B Tree completion illustration on biological trees

#### Data description

In order to demonstrate the tree completion process of the *k*-NCL algorithm, we apply it to two phylogenetic trees of primates, denoted by *T*_1_ and *T*_2_, each defined on partially overlapping sets of taxa (see Figure 5). Let *L*(*T*_1_) and *L*(*T*_2_) be the sets of taxa in *T*_1_ and *T*_2_, respectively.

**Fig. 5.**
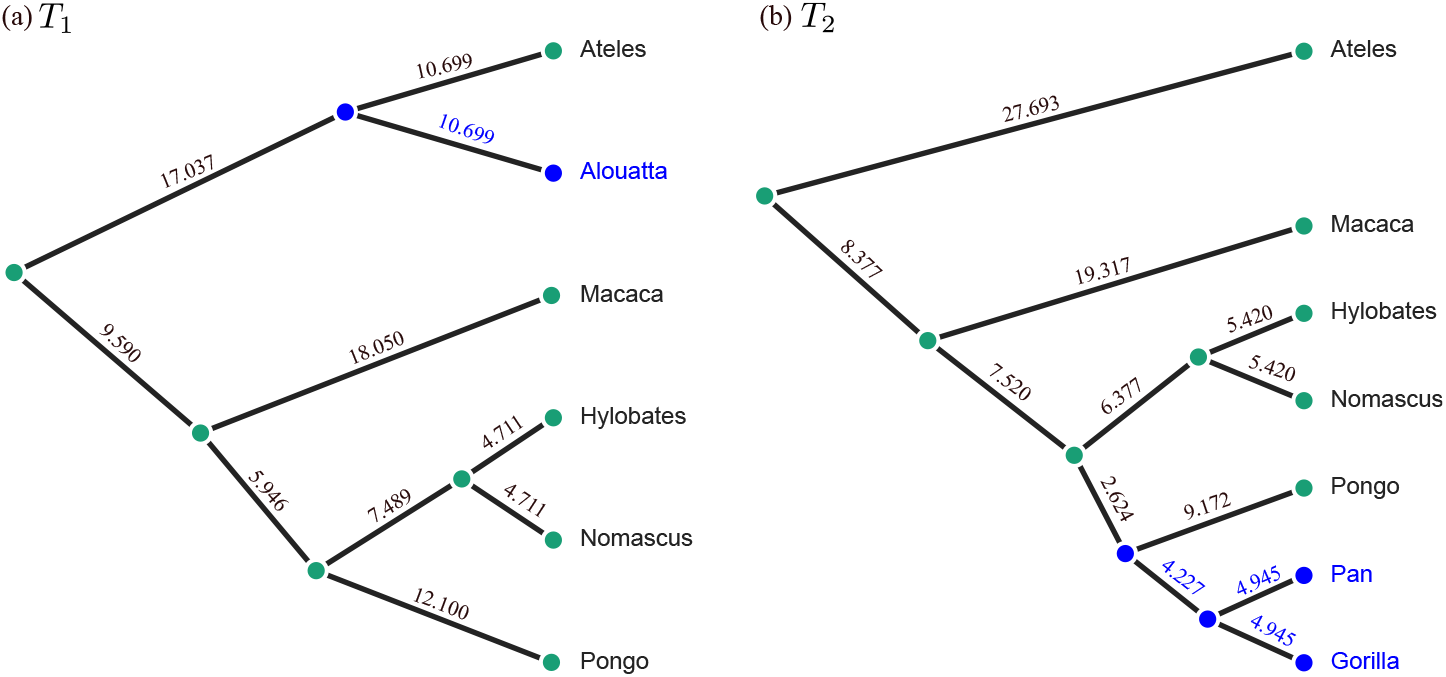
Two phylogenetic trees defined on different but overlapping sets of taxa. (a) Tree *T*_1_ has one distinct leaf (*Alouatta*) and (b) tree *T*_2_ includes two distinct leaves (*Gorilla* and *Pan*). Distinct leaves and their parent nodes are colored blue.

The first tree, *T*_1_, contains six species: *Hylobates, Nomascus, Pongo, Macaca, Ateles*, and *Alouatta*. The second tree, *T*_2_, includes seven species: *Hylobates, Nomascus, Pongo, Macaca, Pan, Gorilla*, and *Ateles*. The two trees share five species: *Hylobates, Nomascus, Pongo, Macaca*, and *Ateles*, denoted by the common leaf set *CL*(*T*_1_, *T*_2_). The total number of unique species across both trees (i.e., *L*(*T*) = *L*(*T*_1_) ∪ *L*(*T*_2_)) is eight.

The goal is to construct two completed trees 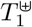 and 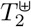 that include all taxa from *L*(*T*) = *L*(*T*_1_) ∪ *L*(*T*_2_). All branch lengths and calculated values in this example are rounded to three decimal places.

#### Tree completion process

Due to the symmetry of the *k*-NCL algorithm (see Proposition 2), the tree completion process can be initiated from either of the input trees, *T*_1_ or *T*_2_, without affecting the final result.

Using Definition 2, the set of common leaves *CL*(*T*_1_, *T*_2_) includes leaves *Hylobates, Nomascus, Pongo, Macaca*, and *Ateles*. The distinct leaves in *T*_1_ and *T*_2_ are *DL*(*T*_1_ | *T*_2_) = {*Alouatta*}, *DL*(*T*_2_ | *T*_1_) = {*Gorilla, Pan*}. In this example, the value of *k* = 3 is chosen. This value is justified by the recommended default value described in the evaluation section. The required parameters for subtree insertion are summarized in Table 1.

#### Completing T_1_ with subtrees from T_2_

To complete tree *T*_1_, the maximal distinct-leaf subtree *S*_2_ = {*Gorilla, Pan*} is identified from tree *T*_2_. This subtree does not share any leaves with *T*_1_, and must be integrated into *T*_1_ to form the completed tree 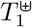. The global adjustment rate *r*(*T*_1_ | *T*_2_) = 0.982 is computed using Equation 1. This rate scales the branch lengths in *S*_2_ to make them compatible with the evolutionary distances in *T*_1_. For subtree *S*_2_, the algorithm selects its three nearest common leaves in *T*_2_, which are *Pongo, Hylobates*, and *Nomascus*, based on branch length proximity. These leaves serve as references for calculating the corresponding leaf-based adjustment rates, which are *r*^(*Pongo*)^(*T*_1_ | *T*_2_) = 0.992, *r*^(*Hylobates*)^(*T*_1_ | *T*_2_) = 0.976, and *r*^(*Nomascus*)^(*T*_1_ | *T*_2_) = 0.976, using Equation 3.

Next, the position distances from each of these three common leaves in *T*_1_ are calculated using Equation 4. These distances are 9.099 for *Pongo*, and 14.075 for both *Hylobates* and *Nomascus*. The candidate insertion point is then selected by minimizing the objective function *f*_*e*_(*x*) (Equation 6). The optimal insertion point, denoted *v*_1_, lies along an internal branch of *T*_1_, where the adjusted subtree *S*_2_ is inserted (Figure 6). The resulting completed tree 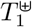 now includes all leaves from both original trees, as shown in Figure 6 (a). This integration preserves the internal relationships within *T*_1_ while extending it with phylogenetic information from *T*_2_.

**Fig. 6.**
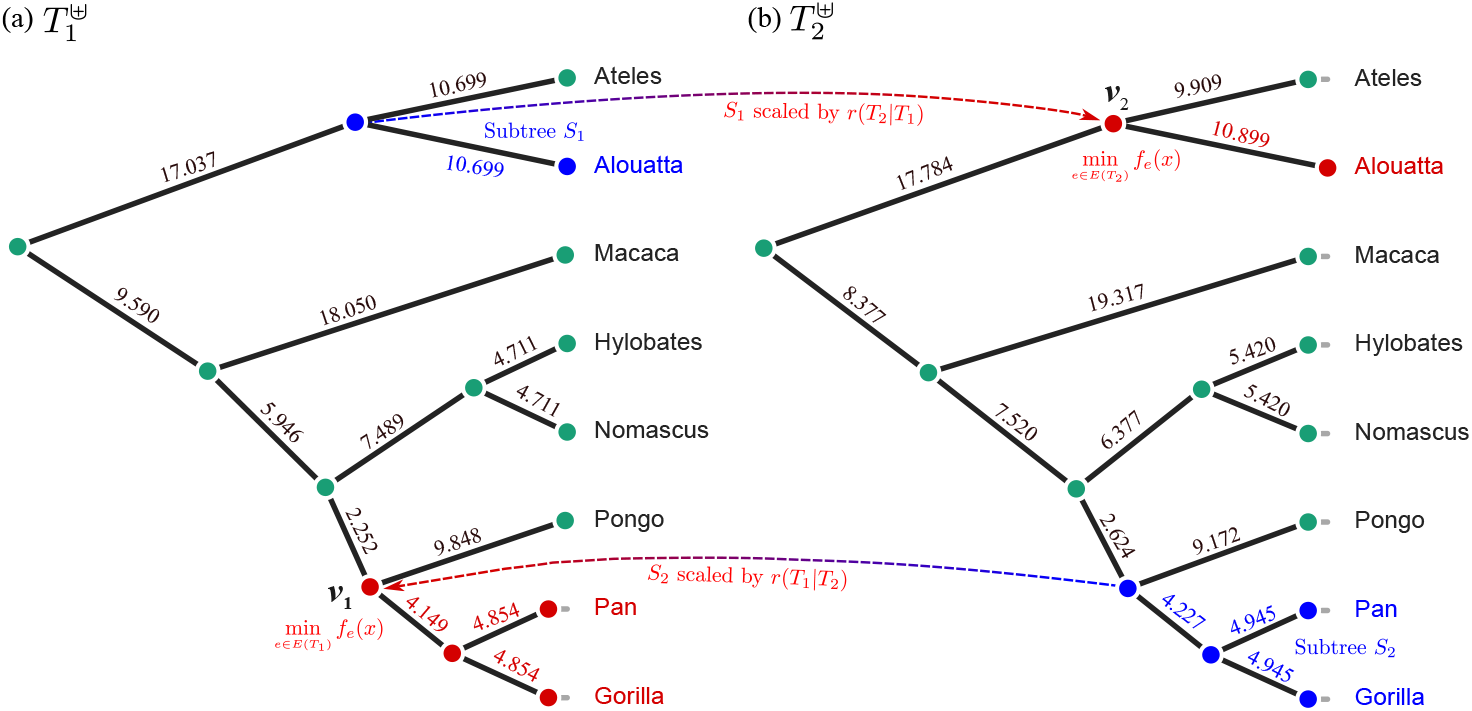
Completed trees 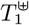 and 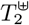. The distinct leaves and their associated internal nodes from the original trees *T*_1_ and *T*_2_ are colored blue. Newly added leaves and their associated internal nodes are colored red. Associated internal nodes comprise all nodes of each distinct-leaf subtree, including their attachment nodes. Arrows indicate the addition of leaves with adjusted branches from one tree to the other.

#### Completing T_2_ with subtrees from T_1_

In the reverse direction, the tree completion process adds the maximal distinct-leaf subtree *S*_1_ = {*Alouatta*}, originally located in *T*_1_, into tree *T*_2_. *Alouatta* is not present in *T*_2_, making it necessary to incorporate it into *T*_2_ to obtain the completed tree 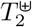. The global adjustment rate *r*(*T*_2_ | *T*_1_) = 1.019 is computed in the same way as before. The *k* nearest common leaves to *S*_1_ in *T*_1_ are identified as *Ateles, Pongo*, and *Macaca*, based on their proximity in the tree structure. For each of these nearest leaves, the corresponding leaf-based adjustment rates are computed, resulting in values *r*^(*Ateles*)^(*T*_2_|*T*_1_) = 0.999, *r*^(*P ongo*)^(*T*_2_|*T*_1_) = 1.008, and *r*^(*Macaca*)^(*T*_2_|*T*_1_) = 1.045.

Based on these values, the position distances from the selected common leaves in *T*_2_ to the expected insertion point of *S*_1_ are calculated. The distances are 10.688 for *Ateles*, 45.030 for *Pongo*, and 46.687 for *Macaca*. These distances are used to locate a possible insertion point in *T*_2_ via the objective function, whose minimum identifies the optimal location *v*_2_ along a specific branch. The subtree *S*_1_, scaled by the adjustment rate *r*(*T*_2_ | *T*_1_), is inserted at this point, as shown in Figure 6 (b). The resulting completed tree 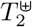 includes all taxa from both *T*_1_ and *T*_2_, preserving the original structure of *T*_2_ while integrating the distinct taxon from *T*_1_.

#### Results of completing T_1_ and T_2_

The outcome of the tree completion process is illustrated in Figure 6, which shows the completed trees 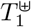 and 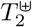 with all taxa integrated.

This example demonstrates how the *k*-NCL algorithm systematically completes phylogenetic trees by integrating distinct-leaf subtrees through distance-based placement and branch length adjustment. The mutual completion of *T*_1_ and *T*_2_ results in two augmented trees defined on the same set of taxa, while preserving the evolutionary structure of the original trees.

### C Supplementary tables

The details of the datasets used in this study are summarized in Table 2.

The cluster separation metrics (silhouette and Dunn) for the tree completion method comparison part are presented in Table 3.

### D Algorithm pseudocode

The pseudocode for the *k*-NCL algorithm is provided below to illustrate the step-by-step procedure underlying the method described in this work.

The *k*-NCL algorithm can be summarized at a high level as follows. Each phylogenetic tree *T*_*i*_ (for *i* ∈ {1, 2}) is completed by integrating distinct leaves from the other tree *T*_3−*i*_. The process begins by identifying the common leaf set *CL*(*T*_1_, *T*_2_), along with the set of distinct leaves *DL*(*T*_3−*i*_ | *T*_*i*_) (Definition 2). Based on this, maximal distinct-leaf subtrees (Definition 3) are extracted. They represent parts of *T*_3−*i*_ that should be inserted into *T*_*i*_. To preserve the input tree, a working copy 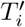 is created as an exact duplicate of *T*_*i*_ (same topology and branch lengths), and all insertions are performed on 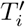, leaving *T*_*i*_ unchanged. The extracted subtrees are scaled using a global adjustment rate (Equation 1) that aligns the average pairwise distances between the common leaves of the two trees. In order to determine where each subtree from *T*_3−*i*_ should be inserted into 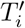, the *k* nearest common leaves (Definition 4) are identified from branch length proximity in *T*_3−*i*_ and then used to calculate position distances in 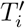 through local scaling factors (Equation 4). An optimal insertion point is selected by minimizing an objective function (Equation 6) that compares position distances to observed distances (Equation 5) in 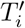. Once the optimal location is found, the subtree is inserted into 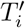 with adjusted branch lengths. This procedure is applied iteratively for all maximal distinct-leaf subtrees from *T*_3−*i*_, resulting in a completed version of *T*_*i*_ that incorporates all leaves from both original trees. The same method is then repeated in the reverse direction to complete *T*_3−*i*_, producing two mutually completed trees, 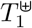 and 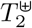.

#### Algorithm 1

*k*-Nearest Common Leaves algorithm

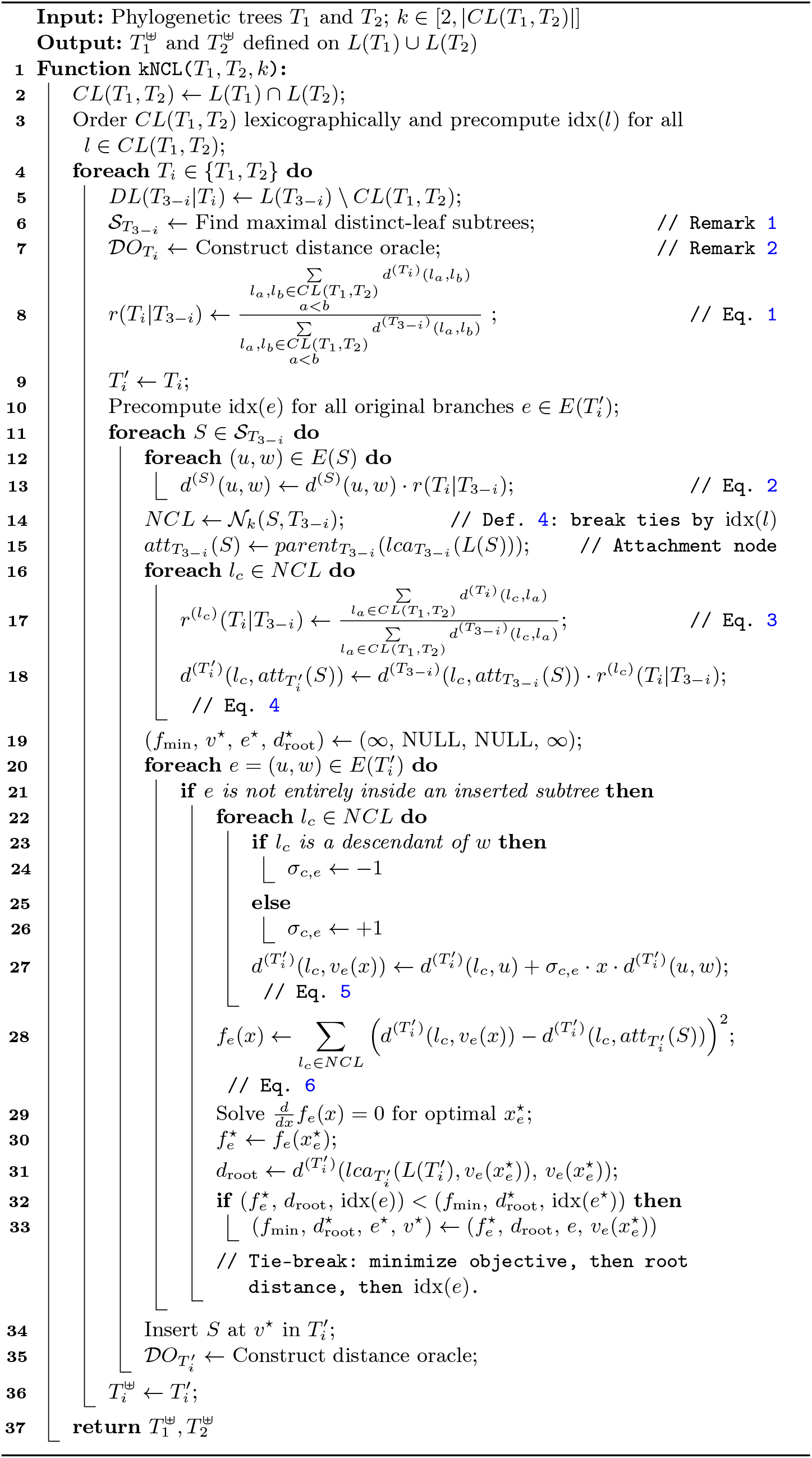

## References

1. Bansal, M.S.: Linear-time algorithms for phylogenetic tree completion under robinson–foulds distance. Algorithms Mol. Biol. 15, 1–15 (2020)

2. Bansal, M.S., Burleigh, J.G., Eulenstein, O., Fernández-Baca, D.: Robinson-foulds supertrees. Algorithms Mol. Biol. 5(1), 1–12 (2010)

3. Bender, M.A., Farach-Colton, M.: The lca problem revisited. In: Latin American Symposium, Punta del Este, Uruguay, April 10-14, 2000 Proceedings 4. pp. 88–94. Springer (2000)

4. Billera, L.J., Holmes, S.P., Vogtmann, K.: Geometry of the space of phylogenetic trees. Adv. Appl. Math. 27(4), 733–767 (2001)

5. Chen, D., Burleigh, J.G., Bansal, M.S., Fernández-Baca, D.: Phylofinder: an in-telligent search engine for phylogenetic tree databases. BMC Evol. Biol. 8, 1–11 (2008)

6. Christensen, S., Molloy, E.K., Vachaspati, P., Warnow, T.: Octal: Optimal completion of gene trees in polynomial time. Algorithms Mol. Biol. 13(1), 1–18 (2018)

7. Cotton, J.A., Wilkinson, M.: Majority-rule supertrees. Syst. Biol. 56(3), 445–452 (2007)

8. Dong, J., Fernández-Baca, D.: Properties of majority-rule supertrees. Syst. Biol. 58(3), 360–367 (2009)

9. Dunn, J.C.: Well-separated clusters and optimal fuzzy partitions. Journal of Cybernetics 4(1), 95–104 (1974)

10. Grindstaff, G., Owen, M.: Geometric comparison of phylogenetic trees with different leaf sets. arXiv preprint 1807.04235 (2018)

11. Hinchliff, C.E., Smith, S.A., Allman, J.F., Burleigh, J.G., Chaudhary, R., Coghill, L.M., Crandall, K.A., Deng, J., Drew, B.T., Gazis, R., et al.: Synthesis of phylogeny and taxonomy into a comprehensive tree of life. PNAS 112(41), 12764– 12769 (2015)

12. Jetz, W., Pyron, R.A.: The interplay of past diversification and evolutionary isolation with present imperilment across the amphibian tree of life. Nat. Ecol. Evol. 2(5), 850–858 (2018)

13. Jetz, W., Thomas, G.H., Joy, J.B., Hartmann, K., Mooers, A.O.: The global diversity of birds in space and time. Nature 491(7424), 444–448 (2012)

14. Koshkarov, A., Tahiri, N.: Novel algorithm for comparing phylogenetic trees with different but overlapping taxa. Symmetry 16(7), 790 (2024)

15. Kupczok, A.: Split-based computation of majority-rule supertrees. BMC Evol. Biol. 11(1), 1–13 (2011)

16. Li, W., Koshkarov, A., Tahiri, N.: Comparison of phylogenetic trees defined on different but mutually overlapping sets of taxa: A review. Ecol. Evol. 14(8), e70054 (2024)

17. Llabrés, M., Rosselló, F., Valiente, G.: The generalized robinson-foulds distance for phylogenetic trees. J. Comput. Biol. 28(12), 1181–1195 (2021)

18. Mahbub, S., Sawmya, S., Saha, A., Reaz, R., Rahman, M.S., Bayzid, M.S.: Quartet based gene tree imputation using deep learning improves phylogenomic analyses despite missing data. J. Comput. Biol. 29(11), 1156–1172 (2022)

19. Mai, U., Mirarab, S.: Completing gene trees without species trees in sub-quadratic time. Bioinformatics 38(6), 1532–1541 (2022)

20. McArthur, R.N., Zehmakan, A.N., Charleston, M.A., Lin, Y., Huttley, G.: Spectral cluster supertree: fast and statistically robust merging of rooted phylogenetic trees. Front. Mol. Biosci. 11, 1432495 (2024)

21. Priel, A., Tamir, B.: A vectorial tree distance measure. Sci. Rep. 12(1), 5256 (2022)

22. Rabiee, M., Mirarab, S.: Instral: discordance-aware phylogenetic placement using quartet scores. Syst. Biol. 69(2), 384–391 (2020)

23. Real, R., Vargas, J.M.: The probabilistic basis of jaccard’s index of similarity. Syst. Biol. 45(3), 380–385 (1996)

24. Ren, Y., Zha, S., Bi, J., Sanchez, J.A., Monical, C., Delcourt, M., Guzman, R.K., Davidson, R.: A combinatorial method for connecting bhv spaces representing different numbers of taxa. arXiv preprint 1708.02626 (2017)

25. Robinson, D.F., Foulds, L.R.: Comparison of phylogenetic trees. Math. Biosci. 53, 131–147 (1981)

26. Rousseeuw, P.J.: Silhouettes: a graphical aid to the interpretation and validation of cluster analysis. J. Comput. Appl. Math. 20, 53–65 (1987)

27. Schaller, D., Hellmuth, M., Stadler, P.F.: Asymmetree: A flexible python package for the simulation of complex gene family histories. Software 1(3), 276–298 (2022)

28. Stein, R.W., Mull, C.G., Kuhn, T.S., Aschliman, N.C., Davidson, L.N., Joy, J.B., Smith, G.J., Dulvy, N.K., Mooers, A.O.: Global priorities for conserving the evolutionary history of sharks, rays and chimaeras. Nat. Ecol. Evol. 2(2), 288–298 (2018)

29. Tahiri, N., Fichet, B., Makarenkov, V.: Building alternative consensus trees and supertrees using k-means and robinson and foulds distance. Bioinformatics 38(13), 3367–3376 (2022)

30. Tahiri, N., Willems, M., Makarenkov, V.: A new fast method for inferring multiple consensus trees using k-medoids. BMC Evol. Biol. 18, 1–12 (2018)

31. Thorup, M., Zwick, U.: Approximate distance oracles. J. ACM 52(1), 1–24 (2005)

32. Upham, N.S., Esselstyn, J.A., Jetz, W.: Inferring the mammal tree: species-level sets of phylogenies for questions in ecology, evolution, and conservation. PLoS Biol. 17(12), e3000494 (2019)

33. Wang, J.T., Shan, H., Shasha, D., Piel, W.H.: Fast structural search in phylogenetic databases. Evol. Bioinform. 1, 117693430500100009 (2005)

34. Whidden, C., Zeh, N., Beiko, R.G.: Supertrees based on the subtree prune-andregraft distance. Syst. Biol. 63(4), 566–581 (2014)

35. Yao, K., Bansal, M.S.: Optimal completion and comparison of incomplete phylogenetic trees under robinson-foulds distance. In: 32nd Annual Symposium on Combinatorial Pattern Matching (CPM 2021). Schloss Dagstuhl-Leibniz-Zentrum für Informatik (2021)

36. Yasui, N., Vogiatzis, C., Yoshida, R., Fukumizu, K.: imphy: Imputing phylogenetic trees with missing information using mathematical programming. IEEE/ACM Trans. Comput. Biol. Bioinf. 17(4), 1222–1230 (2018)

37. Yoshida, R.: Imputing phylogenetic trees using tropical polytopes over the space of phylogenetic trees. Mathematics 11(15), 3419 (2023)

